# Characterization of the HLA-DR immunopeptidome of bronchoalveolar lavage cells in patients with newly diagnosed rheumatoid arthritis and healthy current-smoker controls

**DOI:** 10.1101/2024.12.02.626311

**Authors:** Benedikt Zöhrer, Ákos Végvári, Martina Bonatti, Iryna Kolosenko, Nicole Wagner, Antonio Checa, Vijay Joshua, C. Magnus Sköld, Karin Lundberg, Lars Klareskog, Vivianne Malmström, Åsa M. Wheelock

**Affiliations:** Immunology and Respiratory Medicine Unit, Department of Medicine Solna, Center for Molecular Medicine, Karolinska Institutet Stockholm, Sweden; Proteomics Biomedicum, Division of Chemistry I, Department of Medical Biochemistry and Biophysics, Karolinska Institutet, Stockholm, Sweden; Unit of Integrative Metabolomics, Department of Environmental Medicine, Karolinska Institutet, Stockholm Sweden; Department for Respiratory Medicine and Allergy, Karolinska University Hospital, Stockholm, Sweden; Division of Rheumatology, Department of Medicine Solna, Center for Molecular Medicine, Karolinska Institutet, Karolinska University Hospital, Stockholm, Sweden

**Author notes:** **Correspondence should be addressed to:** Åsa M. Wheelock, Immunology and Respiratory Medicine Unit, Dept. of Medicine Solna and Center for Molecular Medicine, Karolinska Institutet, Visionsgatan 18, CMM, L8:02, 17174 Solna, Sweden.

**Keywords:** HLA-DR antigens, rheumatoid arthritis, bronchoalveolar lavage

## Abstract

Evidence suggests that self-tolerance is breached in the lung prior to the clinical onset of rheumatoid arthritis (RA) in the joints. The human leukocyte antigen DR (HLA-DR) shared epitope (SE) represents the strongest genetic risk factor for sero-positive RA. However, to our knowledge, the HLA-DR immunopeptidome of the RA lung and its link to HLA-DR genotype has not been investigated to date.

The objective of this study was to optimize the methods for characterizing the HLA-DR immunopeptidome of lung immune cells and apply it to newly diagnosed RA patients versus current-smoker healthy controls, as well as to investigate the connection with the HLA-DR genotype.

The HLA-DR immunopeptidome method was improved to facilitate characterization from as few as 6 million bronchoalveolar lavage (BAL) cells per subject, consisting primarily of alveolar macrophages. This method was applied to newly diagnosed RA patients naïve to treatment (n=9, LURA cohort), as well as healthy current-smoker controls (n=10, COSMIC cohort). For five of the RA patients, a 6-month follow-up after initiation of the standard-of-care treatment regime was also included. After isolation and purification, peptide samples were separated by nano-flow liquid chromatography coupled to an Orbitrap mass spectrometer equipped with ion mobility device (FAIMS). Mass spectra acquired in data dependent acquisition mode were then searched against a human proteome database. Subsequently, the identified peptides were deconvoluted to their predicted binding HLA-DR allele using MHCMotifDecon based on the sequenced genotype of the individual.

An optimized sample preparation and analytic method enabled the detection of over 23,000 peptides from over 3,000 source proteins with between 1,000 and 5,000 peptides identified per sample. Notably, the application of FAIMS with three compensation voltages allowed for efficient transfer of 2+, 3+, and 4+ peptide ions while removing singly charged background ions. Hierarchical clustering revealed that the immunopeptidome was more driven by the HLA-DR genotype than by RA disease or sex. However, since the HLA-DR genotype is a strong risk factor for RA, these results are convoluted. When deconvoluting the peptides to their predicted binding allele, the HLA-DRB1 alleles *01:01, *04:01, *04:04, *04:05, *04:07, and *10:01 were consistently assigned more peptides than other alleles. Except for *04:07 these alleles belong to the SE risk factor alleles, providing a potential explanation between HLA-SE and RA pathogenesis. Native peptides from known citrullinated and non-modified RA autoantigens (such as α-enolase and calreticulin) were detected and validated as binders in prediction algorithms. No significant differences were found between base line and follow-up (post-treatment) samples from RA patients.

Taken together, this data characterizes the HLA-DR immunopeptidome in the lung of early RA in an unprecedented manner, which together with future immunogenicity studies will help our understanding of the connection between the lung and the pathogenesis of RA. Finally, more peptides predicted to bind to SE alleles and *04:07 compared to other alleles demands further study on the relative expression of HLA-DR alleles and presentation mechanisms to understand the implications for RA.

## 1. Introduction

Rheumatoid arthritis (RA) is a chronic, inflammatory auto-immune disease affecting the joints (1). Currently, 0.5-1.0% of the global population are affected by RA and the available therapies are only able to reduce the symptoms without curing the underlying disease (1, 2). Characteristic for RA is the presence of autoantibodies, which target self-proteins carrying the post-translational modification citrulline, termed anti-citrulline protein antibodies (ACPA)s. These autoantibodies are present in 2/3 of RA patients, often years before disease onset, and are associated with more severe disease trajectory (3). The strongest genetic predisposition for developing ACPA-positive RA is associated with a cluster of polymorphisms within the human leukocyte antigen DR protein (HLA-DR), commonly referred to as the “shared epitope” alleles (SE) (4). The SE is located on the third hypervariable region of HLA-DRB1 in amino acid positions 70-74, and include the sequences QKRAA (e.g., *04:01), QRRAA (e.g., 01:01, *04:04, *04:05, *04:08, and *14:02), and RRRAA (e.g., 10:01). Non-SE alleles can have the motifs DERAA, QRRAE, RRRAE, DRRAA, QARAA, and QKRGR. The HLA-DR molecule plays a crucial role in antigen presentation to CD4+ T cell receptors (TCRs), thereby serving as a key facilitator in the initiation of a robust humoral immune response. However, the mechanistic relationship between HLA-DR SE, the formation of ACPAs, and the pathogenesis of sero-positive RA is not yet understood.

The detection of ACPAs in serum before their appearance in synovial fluid suggests that the immune tolerance towards citrullinated proteins is not breached in the joints (5, 6). Instead, an increasing body of evidence points towards the lung as the organ where the anti-citrullinated autoreactivity first occurs: Smoking, a major environmental risk factor for developing RA, has been shown to increase the citrullination in the lung (7). Furthermore, the lungs of early RA patients shows structural changes (8). In addition, highly mutated ACPA expressing B cells have been identified in the lung of early RA patients and at-risk individuals (9). Yet to our knowledge, the HLA-DR antigen presentation landscape in the RA lung has not been investigated to date. This can be achieved using mass-spectrometry based HLA-DR immunopeptidomics. This technique was previously applied on synovial macrophages and synovial fluid pulsed PBMC-derived dendritic cells from RA patients (10–13). The detected HLA-DR ligands were then tested for immunogenicity in T cell activation assays, identifying known and novel autoantigens.

In the present study, we investigated the HLA-DR immunopeptidomics landscape in the lung of early RA patients naïve to treatment, and age-matched current-smoker controls using our optimized immunopeptidome method that facilitates analysis of scarce samples. By predicting the HLA-DR binding partner for each peptide we are able investigate SE associated peptides especially in heterozygous individuals (14). From the resulting dataset we aim to get insights into the HLA-DR antigen presentation landscape in RA and find potential pathogenic mechanisms for RA.

## 2. Methods

### 2.1 Bronchoalveolar lavage samples from RA and controls

Immune cells from the lung obtained by bronchoalveolar lavage (BAL) were collected from two distinct cohorts recruited at the Karolinska University Hospital: Nine subjects with rheumatoid arthritis were drawn from the LUng investigation in early Rheumatoid Arthritis (LURA) cohort (8). Those individuals were newly diagnosed with RA according to the American College of Rheumatology 1987 classification criteria and had not received prior treatment with oral glucocorticoids or disease modifying antirheumatic drugs (DMARDS). Exclusion criteria included pregnancy, as well as alcohol and drug abuse. Additionally, a subset of five subjects underwent a 6-month follow-up bronchoscopy after initiation of the treatment regimen. The control group consisting of ten smokers with normal lung function were selected from the Karolinska COSMIC (Clinical & Systems Medicine Investigations of Smoking-related Chronic Obstructive Pulmonary Disease) cohort (https://clinicaltrials.gov/study/NCT02627872) (15). The selection of controls aimed to match for age, to achieve equal sex distribution, and to cover as many HLA-DRB1 alleles as possible, prioritizing those present in the RA group. All control subjects had lung function FEV_1_/FVC>0.70 and did not use any inhaled or oral glucocorticoids. Bronchoscopies from both cohorts were conducted at the same clinic by the same clinician during the same time intervall, following a uniform protocol (16, 17). Cells were stored at -80°C either as cell pellets (LURA) or in freezing medium (COSMIC) until use. Detailed cohort characteristics are summarized in Table 1.

**Table 1.** Cohort characteristics of individuals with RA and controls included in this study.

|  | RA Baseline | RA Follow up | Controls |
| --- | --- | --- | --- |
| n (female/male) | 9 (2/7) | 5 (0/5) | 10 (5/5) |
| Age, median (range) | 60 (43-76) | 63 (43-77) | 56 (48-64) |
| Smoking, current/ex | 6/3 | 4/1 | 10/0 |
| HLA-DR SE alleles, 2/1/0 | 3/4/2 | 2/2/1 | 2/5/3 |
| RF positive, n (%) | 9 (100%) | 5 (100%) | N/A |

| ACPA positive, n (%) | 8 (89%) | 4 (80%) | 0 (0%) |
| --- | --- | --- | --- |
| Million BAL cells used, median (range) | 6.0 | 6.0 | 7.1 (6.0-9.0) |
| Macrophages, median (range) | 93% (77-97%) | 93% (93-97%) | 97% (91-98%) |

*Abbreviations: RA – rheumatoid arthritis, SE – Shared epitope, RF – rheumatoid factor, ACPA – anti-citrullinated protein antibody, BAL – bronchoalveolar lavage*

### 2.2 Culturing of Priess cell line (Priess)

Epstein-Barr virus (EBV) transformed B cell line (Priess) was cultured in RPMI medium including L-Glutamine (Gibco, Grand Island, NY, USA) supplemented with 10% (v/v) fetal bovine serum. Cells were washed twice with PBS buffer (Gibco), counted, pelleted in aliquots of 10^7^ cells, and frozen at -80°C until use.

### 2.3 Cross linking of anti-HLA-DR antibody to sepharose protein A beads

5 mL Protein A Sepharose CL4B beads (Cytiva, Uppsala, Sweden) were suspended in 5 mL 0.2 M sodium borate solution pH 9.0 (boric acid purchased from Sigma, Burlington, MA, USA). 12.5 g mouse anti-human HLA-DR antibody (clone L243) were added, and the beads were gently shaken for 1 hour at room temperature. The antibody was kindly provided by prof. Rikard Holmdahl. After binding, the beads were washed with 2 x 50 mL 0.2 M sodium borate solution pH 9.0 and resuspended in 10 mL of the same solution. 40 mL of freshly prepared 20 mM dimethylpimelimidate (Sigma) in 0.2 M sodium borate solution pH 9.0 was added and shaken for 30 minutes at room temperature. Subsequently, the solution was discarded, and the beads were washed with 5 mL 0.2 M sodium borate solution pH 9.0 and resuspended in 10 mL 0.2 M ethanolamine solution, pH 8.0 (ethanolamine purchased from Sigma). To remove potentially non-cross-linked antibody, the beads were washed with 25 mL glycine solution, pH 2.7. The beads were then washed with PBS until neutral pH and resuspended in 5 mL PBS with 0.1% (w/v) sodium azide.

### 2.4 HLA-DR peptide isolation

Randomized BAL cell samples were prepared in two batches of 12 samples on two consecutive days, each with two 5 million cell aliquots of the EBV-transformed B cell line Priess as quality control. Cell pellets were suspended in 30 µL of an aequous solution containing Tris (50 mM, GE Healthcare, Chicaco, IL, USA), NaCl (150 mM, Thermo Scientific, Waltham, MA, USA), EDTA (5 mM), and protease inhibitor (Complete mini, Roche, Basel, Switzerland). 30 µL of a lysis solution containing Tris (50 mM), NaCl (150 mM), EDTA (5 mM), and protease inhibitor and 1% (w/v) Zwittergent (N-Dodecyl-N,N-dimethyl-3-ammonio-1-propanesulfonate, Sigma) were added, and samples were shaken for 2 hours on a horizontal shaker (4°C, 200 rpm). After lysis, the samples were centrifugated for 1 hour (21,100xg, 4°C).

The supernatant was passed through a column containing 30 µL Protein A Sepharose CL4B beads (Cytiva) to remove antibodies. The flowthrough was applied five times to a column containing 75 µg anti HLA-DR antibody (mouse IgG2a clone L243) cross-linked to 30 µL Protein A Sepharose CL4B beads. Next, the column bed was washed three times with each 200 µL Tris (50 mM), NaCl (150 mM), EDTA (5 mM), and protease inhibitor and 0.5% (w/v) Zwittergent to remove non-bound protein, with three times 200 µL Tris (20 mM), NaCl (120 mM) pH 8.0 to remove detergent, once 100 µL Tris (20 mM), NaCl (1 M) pH 8.0, to remove non-specifically binding proteins, again three times 200 µL Tris (20 mM), NaCl (120 mM) pH 8.0, and finally once 100 µL Tris (20 mM) pH 8.0 to remove salt. The column was eluted with 150 µL 1% (v/v) TFA (Sigma) directly onto a column containing 5 mg Oasis HLB particles (Waters, Milford, MA, USA). After washing three times with 0.1% (v/v) TFA, peptides were eluted with 50 µL 35% acetonitrile (Merck, Darmstadt, Germany), 0.1% TFA (v/v) into a Protein LoBind tube (Eppendorf, Hamburg Germany). Samples were dried in a vacuum concentrator (Scanvac Coolsafe, Labogene, Lillerød, Denmark) and stored at -20°C. On the day of the first mass spectrometric analysis, samples were resuspended in 10 µL 2% acetonitrile in water with 0.1% FA (v/v), centrifuged, and transferred to an LC vial.

### 2.5 Mass spectrometric peptide analysis

The isolated HLA-DR immunopeptide samples were analyzed in randomized order in two replicate injections of 5 µL on two separate days, with the second replicate being injected in inverse order compared to the first. During both replicate analyses, 4 and 7 quality control samples respectively were run alongside the BAL cell samples, consisting of isolated HLA-DR peptides from 5 million EBV-transformed B cells (Priess cell line). All samples were analyzed in by nanoLC-MS/MS using an NanoUltimate 3000 coupled to an Orbitrap Eclipse Tribrid mass spectrometer (Thermo) which was equipped with a high-Field Assymetric-waveform Ion Mobility Spectrometry (FAIMS, Thermo). Peptides were trapped on a µPAC trap column (Pharmafluidics, Gent, Belgium), and separated on a 15 cm long TS C18 chromatographic column (IonOpticks, Fitzroy, Australia) using a flow rate of 0.3 µL/min and a column temperature of 35°C. The separation was achieved using a 115 min gradient with eluent A (2% acetonitrile, 0.1% formic acid) and eluent B (98% acetonitrile, 0.1% formic acid) of 0-4% eluent B over 5 min, 4-20% over 70 min, 20-36% over 20 min, 36-95% over 5 min, followed by 5 min isocratic at 95% eluent B before equilibrating for 10 min at 0% eluent B. The FAIMS compensation voltage alternated between –40, –50, and –70 V for 1s each. The mass spectrometer was operated in data-dependent mode with an MS1 precursor selection range of m/z 550-1500, 300-750, and 400-900 for the three compensation voltages respectively. The orbitrap resolution was 120,000 for MS1 and 30,000 for MS2, the MS1 isolation window was set for 0.7, with 30 s dynamic exclusion enabled. HCD fragmentation was performed with a collision energy of 30%.

### 2.6 Peptide search

Peptides were identified from the mass spectrometry raw data using database search tool using MSFragger (v4.1) (18–23). A human protein sequence database (SwissProt, homo sapiens, with the following HLA-DRB allele sequences manually added: B1*01:01, B1*01:02, B1*01:03, B1*03:01, B1*03:02, B1*04:01, B1*04:02, B1*04:03, B1*04:04, B1*04:05, B1*04:07, B1*04:08, B1*07:01, B1*08:01, B1*08:03, B1*08:04, B1*09:01, B1*10:01, B1*11:01, B1*11:03, B1*11:04, B1*12:01, B1*13:01, B1*13:02, B1*13:03, B1*13:05, B1*14:01, B1*14:04, B1*14:06, B1*14:54, B1*15:01, B1*15:02, B1*15:03, B1*16:01, B1*16:02, B3*01:01, B3*02:02, B3*03:01, B4*01:01, B4*01:03, B5*01:01, B5*01:02, B5*02:02) was searched for peptides without protease-specificity with a length between 7 and 25 amino acids. Mass calibration and parameter optimization was enabled, and methionine oxidation (maximum 3 per peptide), N-terminal acetylation, and cysteinylation (both maximum 1 per peptide) were added as variable modifications. Rescoring with MSBooster, PSM validation with Percolator, and protein inference using ProteinProphet were enabled, FDR of 1% was applied on an ion, PSM, and peptide level using Philosopher.

In a separate search, citrullination of arginine and deamidation of asparagine and glutamine were added as variable modifications. For this search, MSBooster was disabled.

A separate search of the proteomes of common lung microbiota was performed, including Apergillus, Streptococcus pneumoniae/pyogenes, Staphylococcus aureus, Pseudomonas aeruginosa, Haemophilus influenzae, Veillonella parvula, Prevotella, and Neisseria).

### 2.7 Genotyping

HLA-DRB1, DRB3, DRB4, and DRB5 genotypes were determined by whole exome sequencing (Novogene, Cambridge, UK). DNA was extracted from PBMC of the subjects and sequenced using the NovaSeq X Plus Series (PE150) with a sequencing depth of 12 G per sample. Subsequently, the genotypes were extracted using Novogene’s in house bioinformatics pipeline.

### 2.8 HLA-DR binding prediction and motif deconvolution

Identified peptides were assigned to their predicted binding HLA-DRB1, and where applicable DRB3, DRB4, and DRB5 alleles in each sample using MHCMotifDecon algorithm (v1.1) and the HLA-DR genotypes. Peptides between 9 to 25 amino acids were included, and a binding rank score of 20% was set as a threshold to filter out non-binding peptides (14).

### 2.9 Testing for ACPA

Study subjects were tested in duplicates for plasma ACPA using the IMMUNOSCAN CCPlus assay (SVAR Life Science AB, Malmö, Sweden) according to the manufacturer’s instructions. ACPA positivity of a sample was defined as a response of ≥25 U/mL.

## 3. Results and Discussion

### 3.1. Data inspection for quality control

With an optimized HLA-DR immunopeptidomics workflow and two replicate injections per sample 23.759 peptides were identified deriving from 3044 human proteins in 24 samples, with 1,118-5,240 peptides identified per sample of 6 million BAL cells (Figure 1 A). In particular, the use of the ion mobility device FAIMS with three compensation voltages (-40, -50, and -70) allowed for efficient transfer of 2+, 3+, and 4+ charged peptide ions to the mass spectrometer. In comparison, a study on HLA-II immunopeptides using FAIMS with only two compensation voltages (-50 and -70) showed a decrease in doubly charged peptide spectrum matches (PSMs) of >20% compared to another ion mobility instrumentation and our data (37142057). The use of FAIMS using three compensation voltages removed singly charged mostly non-peptidic background ions while ensuring the detection of peptides spanning the full charge distribution of HLA-DR immunopeptides.

**Figure 1:**
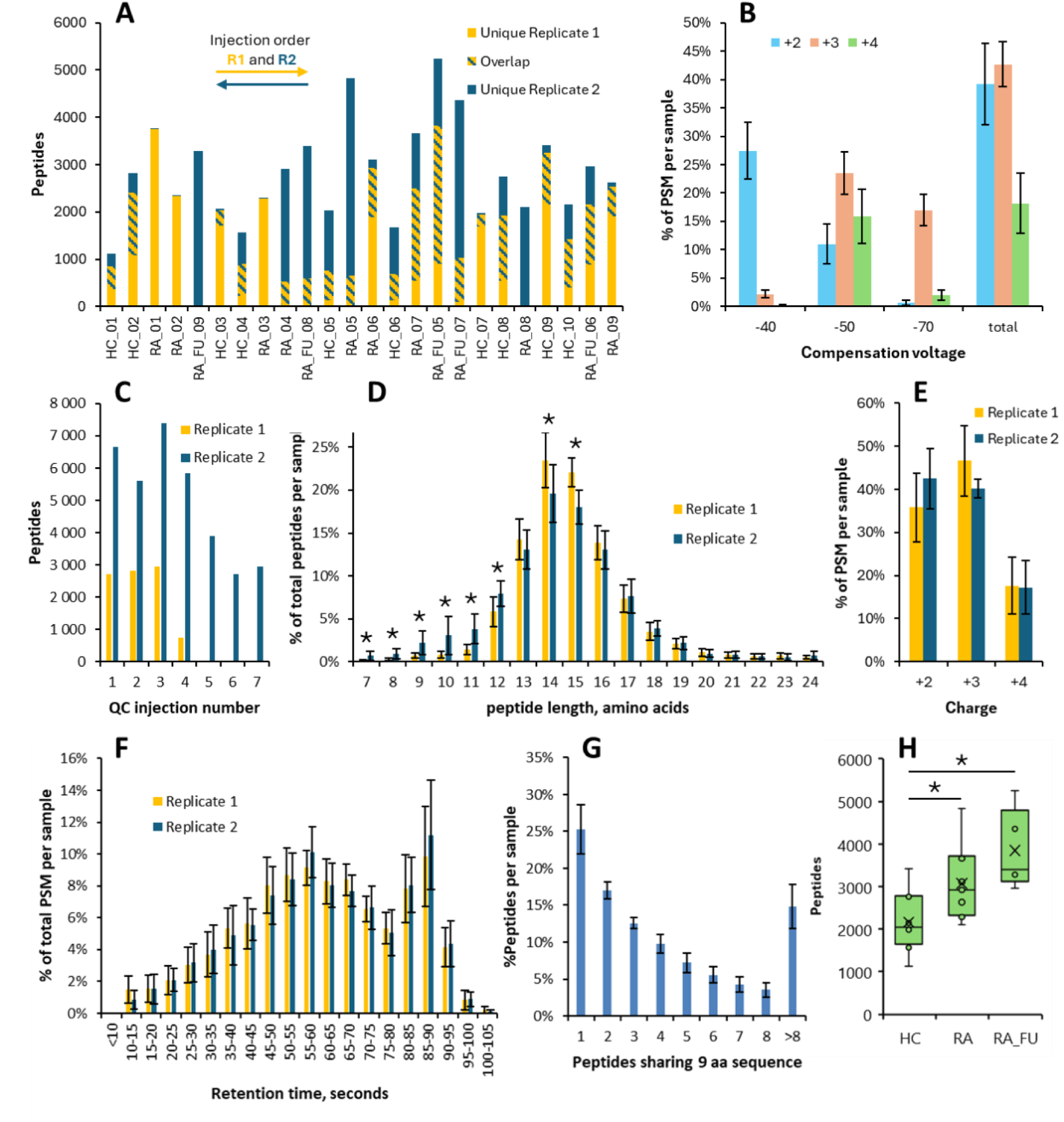
General data characteristics used for quality control. Panel A displays peptides identified exclusively in replicate injection 1 (yellow) or 2 (blue) or in both injections (striped). Arrows indicate the injection order for the respective replicate injections. Average charge distribution of the peptide spectrum matches (PSMs) per sample depending on the compensation voltage of the ion mobility interface is depicted in B, with error bars indicating the standard deviation. Peptide identifications for each quality control (QC) sample measured during injection replicate 1 (n=4) and 2 (n=7) are shown in C. The peptide length distribution (D), PSM charge distribution (E), and elution profile of PSM (F) in the study samples measured in replicate 1 (yellow) and 2 (blue) are shown as average with error bars indicating the standard deviation. Panel G shows which average percentage of peptides per sample share a 9 amino acid long partial sequence with how many peptides in the same sample. Error bars indicate the standard deviation. Panel H shows a boxplot comparing the number of peptide identifications per sample between the groups. HC: Healthy smoker control, RA: RA naïve to treatment, RA_FU: RA following 6 months of pharmacological intervention. Significant differences (p<0.05) between replicate injections (D, E, F) or groups (H) are indicated with an asterisk.

Using dual injections per replicate, we could steeply increase the number of identified peptides per sample and therefore the depth of the dataset (Figure 1 A). The overlap of peptide identifications between any two quality control injections with similar number of peptides (QC1-3 and QC1-4 in replicate injections 1 and 2 respectively, see Figure 1 C) was 54%-62% and 64%-69% for replicate injections 1 and 2 respectively (data not shown). This is expected for immunopeptidomics experiments using DDA (24). Lower reproducibility compared to proteomics analyses on tryptic peptides are method intrinsic and likely associated with less ideal peptide properties for mass spectrometric identification. The number of identified peptides between replicate injections was highly variable and the peptide number in the quality control samples dropped within both replicate runs. Furthermore, one chromatographic column had to be exchanged due to clogging. Challenges with robustness are also common for immunopeptidomics analyses on scarce samples, likely caused by co-isolated contaminants. The reduction of those contaminants by means of additional peptide purifications steps would drastically decrease the peptide yield and result in fewer identified peptides. To ensure that the variability in peptide detection did not cause any bias in the peptide identification, we compared both replicate injections regarding peptide length distribution, PSM charge distribution, and elution profile (Figure 1 D, E, F). While charge and retention time distribution were comparable, the second replicate injection yielded a significantly higher proportion of peptide identifications in the range of 9-12 amino acids. However, since both length distributions are within expectation for HLA-DR immunopeptidomics (gaussian distribution between 9 and 25 amino acids with a maximum at 14-15 amino acids), both replicate datasets were considered acceptable. Additional evidence of a dataset highly enriched with MHCII immunopeptides is the high percentage of peptides (75%) which share a 9 amino acid long sequence, while differing in length (Figure 1 G). This is characteristic, as the peptides are trimmed by endolysosomal peptidases after binding to the MHCII complex, resulting in so called nested sets of peptides.

The number of peptides identified between replicate injections was highly variable, peptide identifications in the quality control samples dropped within one analytical run (Figure 1 C), and one analytical column had to be exchanged due to clogging. From this data we conclude that the data from both replicate measurements can be used. Therefore, we proceed with a Boolean data format, where presence of a peptide in at least one of the replicate injections per sample is considered a positive identification, whereas the absence in both replicates is considered a non-detection. However, due to the scarce clinical material (6 million cells per sample), and the data dependent nature of the analysis, non-detection does not necessarily signify absence of the peptide. This needs to be considered for all subsequent analyses.

When comparing the groups, both the treatment naïve and post-treatment RA groups displayed significantly more peptide identifications per sample than the healthy smoker controls. Due to the high variability in peptide identifications for the individual measurements and the possibility of other biases (as the RA group and the control groups were collected in distinct cohorts) further data inspection is necessary to formulate a viable hypothesis for these differences.

### 3.2 HLA-DR genotype drives the similarity of the HLA-DR immunopeptidome

To understand which factors were most strongly influencing the HLA-DR immunopeptidome on cells from BAL, we performed hierarchical clustering using Jaccard distance between the samples (Figure 2). Jaccard distance is appropriate for Boolean data for which the presence of a variable is more meaningful than its absence – as it is the case with DDA. As expected, samples from the same individual, i.e. baseline and follow-up pairs of subjects with RA, are most similar (Figure 2). However, the samples do not cluster according to disease state, sex, or ACPA positivity. This indicates that despite the difference in total peptide count between the groups (Figure 1 H) disease- or treatment status does not have primary impact on the data. In contrast, the HLA-DRB1 genotype seems to have a stronger influence on immunopeptidome similarity between the samples. In particular, samples with HLA-DRB1*04:01 or *04:05 genotype display a higher correlation. Interestingly, those are the most frequent risk alleles in population with European and East Asian ancestry, respectively (25, 26). Conversely, the nine samples carrying the *03:01 genotype do not cluster together.

**Figure 2:**
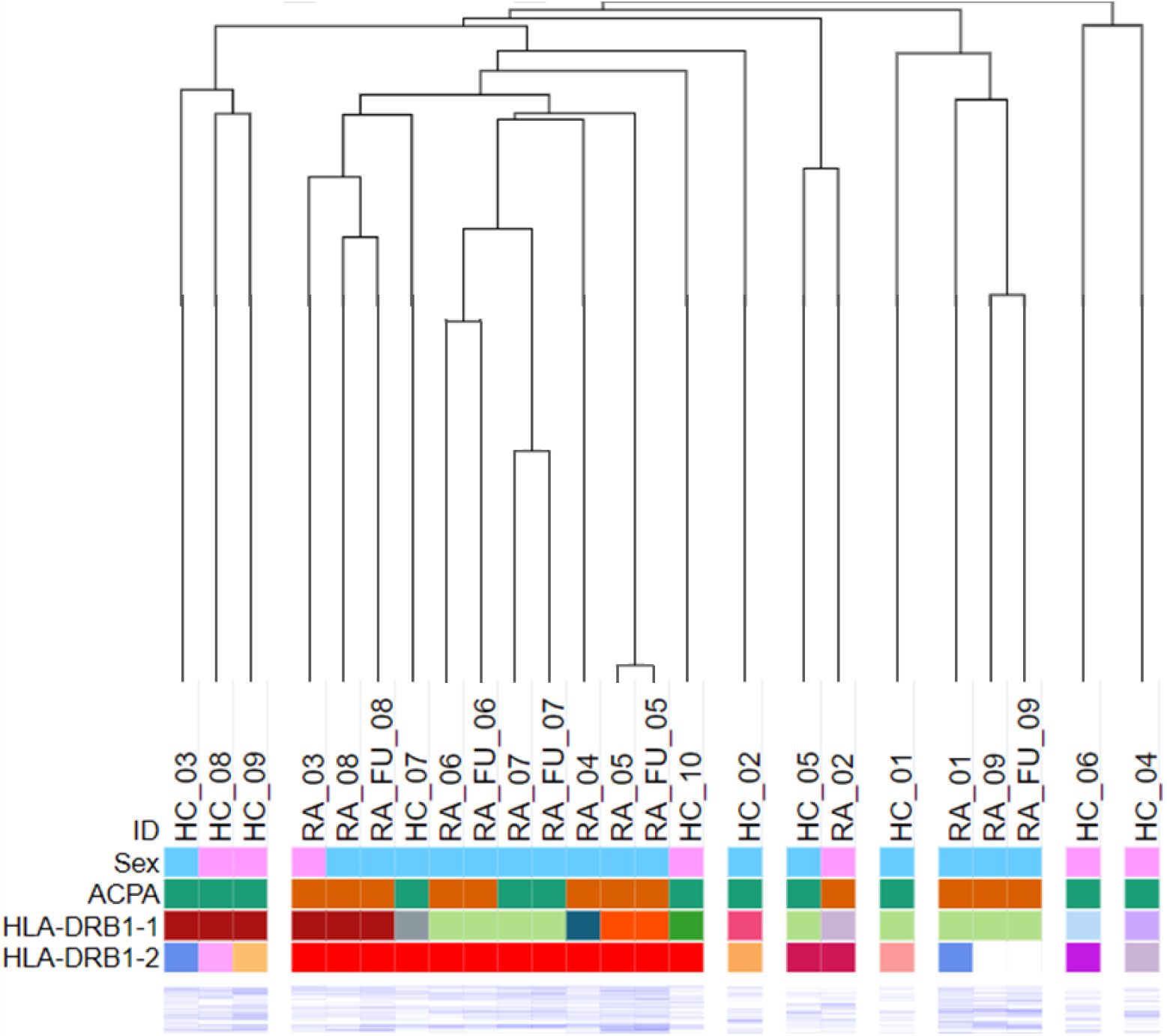
Similarity of the HLA-DR immunopeptidome between the samples. Dendrogram of all identified peptides per sample using Jaccard distance. The peptide heat map has been cropped to show only the first few rows. Figure generated using Morpheus, https://software.broadinstitute.org/morpheus.

### 3.3 More peptides predicted to bind to SE alleles and HLA-DRB1*04:07

Since the HLA-DRB1 genotype best explained peptide similarity between samples, we can assume that the genotype is a stronger driver on the HLA-DR immunopeptidome in the lung than sex, RA diagnosis, or treatment. This observation has been formulated in another cohort investigating the HLA-DR immunopeptidome of various cell types in the context of multiple sclerosis (27). For this reason, we deconvoluted the HLA-DR immunopeptidome of each sample to the constituent HLA-DRB1 genotypes, and where applicable to the genotype of the paralogues DRB3, DRB4, and DRB5. As expected, HLA-DRB3, DRB4, and DRB5 occur according to the ancestral haplotypes, e.g. HLA-DRB1*03:01 and HLA-DRB1*04:01 likely expressing also HLA-DRB3 and DRB4 respectively (Supplementary figure 1 A). The relative and absolute contribution of peptides predicted to bind to DRB4 is lower compared to DRB3, which has been previously reported (Supplementary figure 1B and C) (14).

**Supplementary figure 1.**
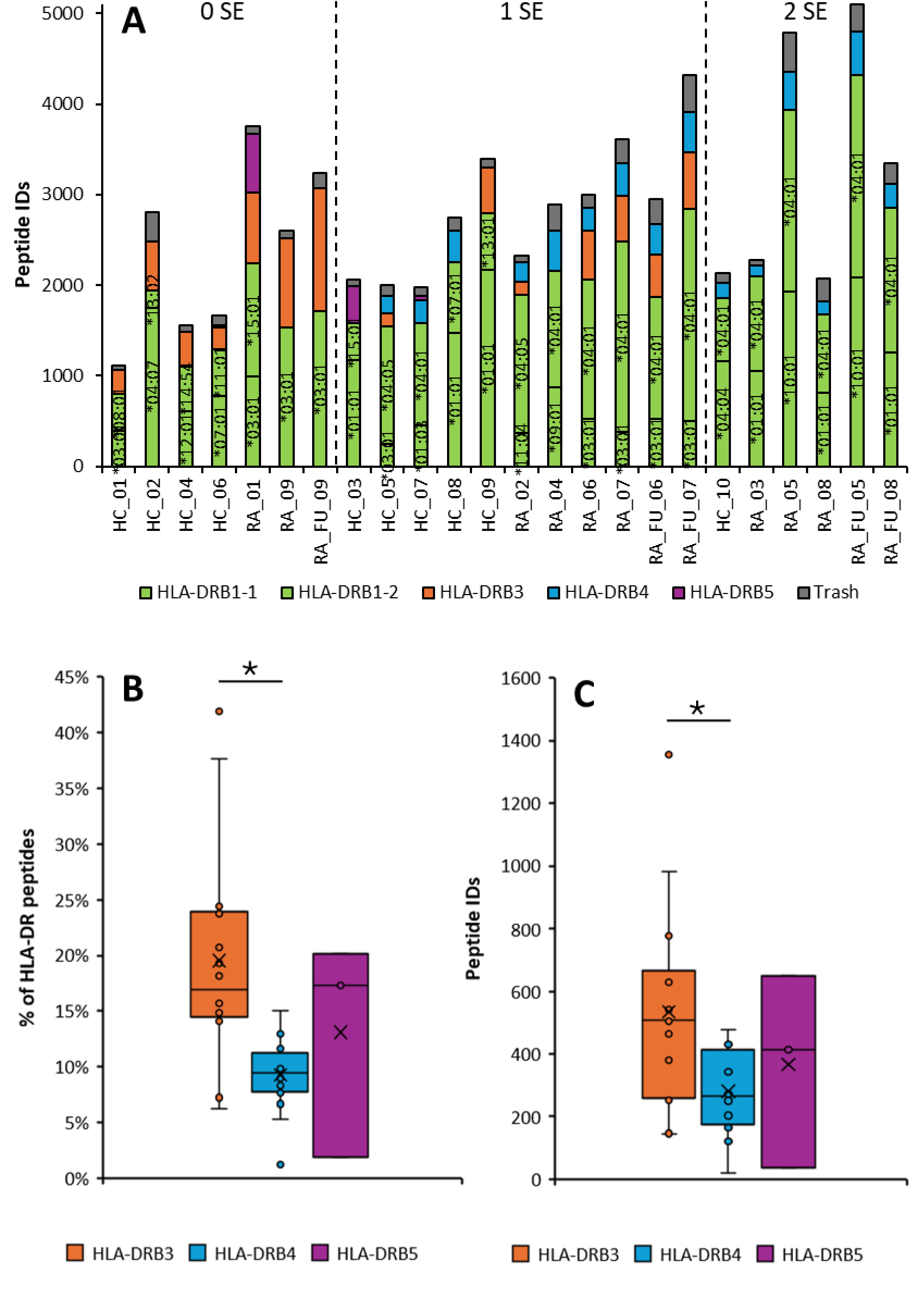
Comparison of peptide numbers deconvoluted to the genotypes of HLA-DRB1, DRB3, DRB4, and DRB5. Panel A shows the number of identified peptides per allele of HLA-DRB1 (green), HLA-DRB3 (orange), DRB4 (blue), and DRB5 (violet). Peptides not predicted to bind (trash) are highlighted in grey. Panel B shows the relative contribution of DRB3, DRB4, DRB5 to the HLA-DR immunopeptidome, while Panel C shows the same peptide numbers. The box in box-and-whiskers-plots represents the interquartile range (IQR), with the lower and upper bounds corresponding to the first and third quartiles. The horizontal line inside the box indicates the median, while the ‘x’ denotes the mean. The whiskers extend to the full range of the data, showing the minimum and maximum values.

When regarding solely HLA-DRB1 peptides, more peptides are predicted to bind on SE alleles compared to non-SE alleles both in relative and absolute numbers (Figure 3). The only exception is *04:07 in HC_02, which is co-expressed with *13:02 and predicted to bind 90% of the DRB1 peptides of this sample. *04:07 does not contain the SE motif, however it has almost identical peptide binding preferences compared to SE allele *04:08. This finding could indicate either an increased allele-specific expression or a broader diversity of presented peptides for SE alleles and *04:07. To further investigate if this observation is caused by a bias in any part of our analytical pipeline, we compared retention time profile and the predicted binding strength of the identified peptides (Figure 4 A and B). These analyses show that SE alleles do not have a consistently different retention time profile and binding strength profile compared to non-SE. This in turn suggests that the increased number of peptides predicted to bind to SE alleles and *04:07 is not likely to be caused by a bias in the chromatographic separation or the binding affinity and could indeed reflect differential expression or presentation profiles of the alleles.

**Figure 3.**
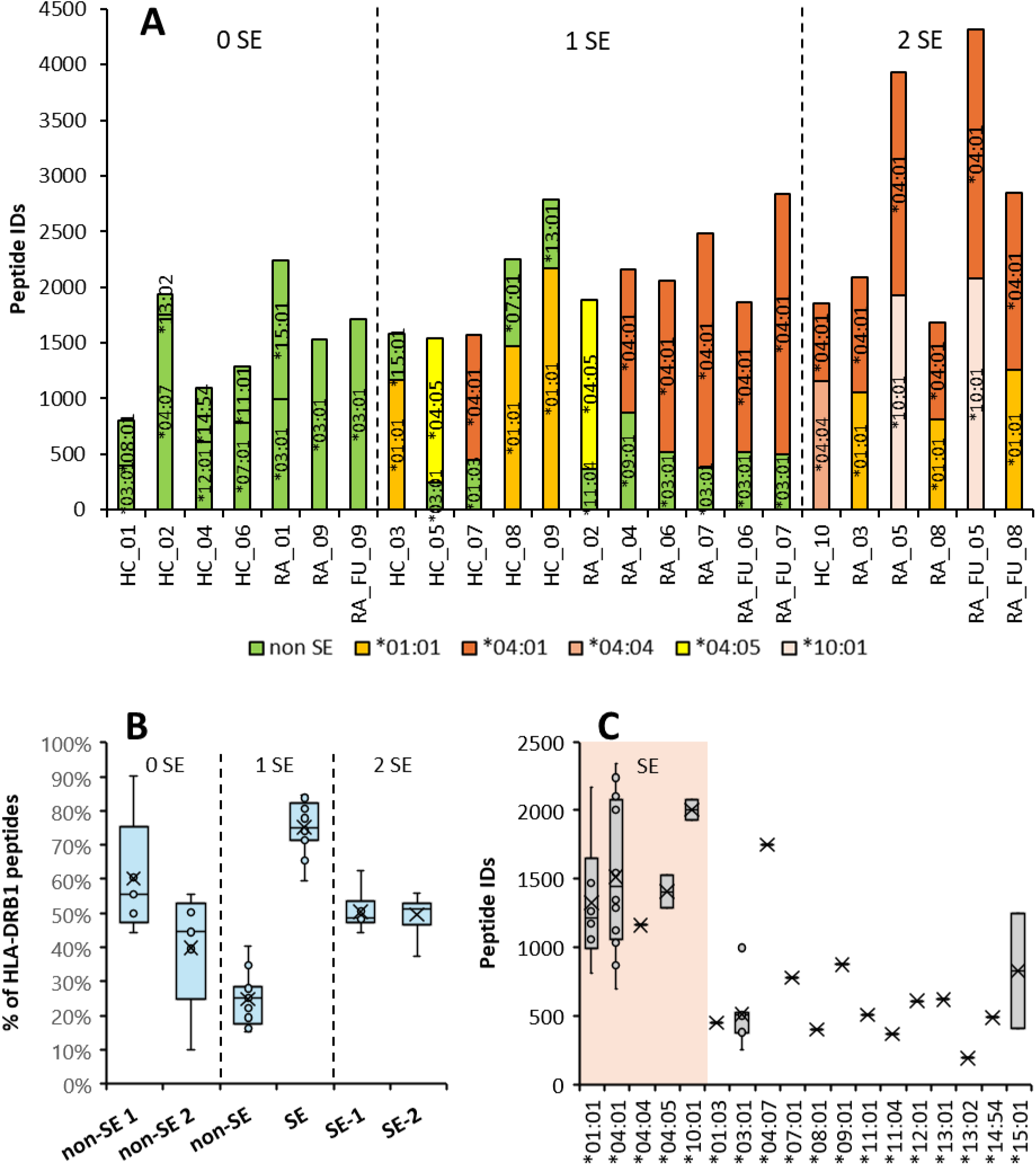
Comparison of peptide numbers deconvoluted to the predicted binding HLA-DRB1 genotypes. Panel A shows the peptide numbers deconvoluted to HLA-DRB1 genotype, with samples grouped by the number of HLA-DR shared epitope (SE) alleles. SE alleles are coded in colors other than green. Panel B summarizes the percentage contribution of SE and non-SE alleles to the HLA-DRB1 peptides, depending on the number of SE alleles per sample. In Panel C the absolute numbers of peptides for each HLA-DRB1 genotype are plotted, with SE genotypes highlighted with an orange background. The box in box-and-whiskers-plots represents the interquartile range (IQR), with the lower and upper bounds corresponding to the first and third quartiles. The horizontal line inside the box indicates the median, while the ‘x’ denotes the mean. The whiskers extend to the full range of the data, showing the minimum and maximum values.

**Figure 4:**
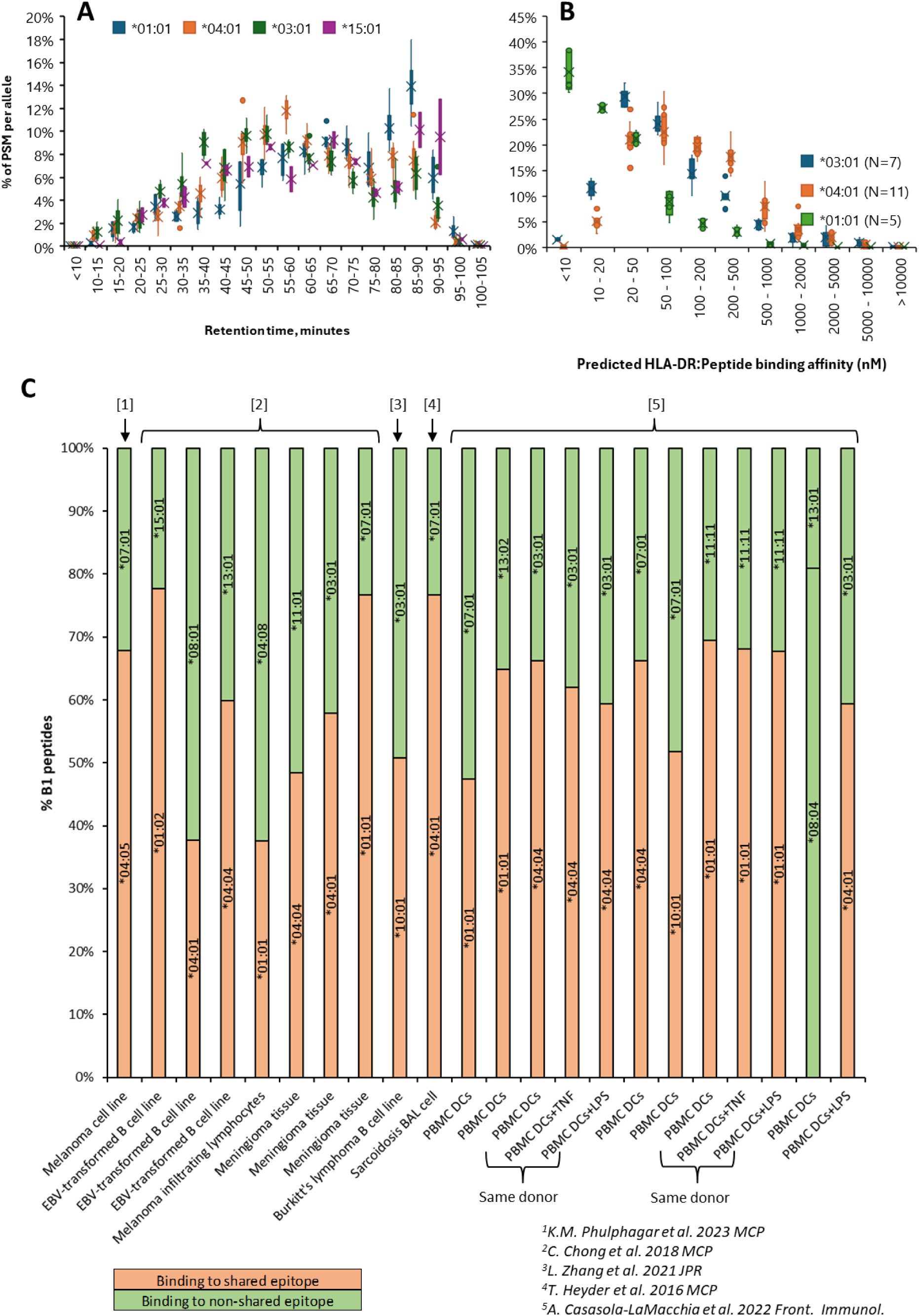
Allele specific peptide properties in the current study and peptide proportions in the literature. Retention time profile for peptides predicted to bind to shared epitope alleles *01:01 (n=6) and *04:01 (n=12) and non-shared epitope alleles *03:01 (n=9) and *15:01 (n=2) normalized to total peptide spectrum matches in each sample predicted to bind to the respective allele. Distribution of predicted binding affinity per allele (*03:01, *04:01, and *01:01) is shown in B. Relative numbers of peptides binging to either HLA-DRB1 allele from deconvoluted publicly available HLA-DR immunopeptidomics datasets are shown in C. The box in box-and-whiskers-plots represents the interquartile range (IQR), with the lower and upper bounds corresponding to the first and third quartiles. The horizontal line inside the box indicates the median, while the ‘x’ denotes the mean. The whiskers extend to the full range of the data, showing the minimum and maximum values.

Next, we investigated publicly available datasets on HLA-DR immunopeptidomics and applied the same peptide deconvolution workflow (MHCMotifDecon) (Figure 4 C). In a BAL cell sample from a smoking subject with sarcoidosis from a previous publication of our laboratory around three times as many peptides were predicted to bind to *04:01 compared to *03:01, which agrees with our current findings (28). Also, in other datasets more peptides were predicted to bind to a SE allele compared to non-SE allele. However, there are exceptions. For example, more peptides are predicted to bind to non-SE allele *08:01 compared to SE allele *04:01 in an EBV-transformed B cell line (29).

Transcriptomics studies also suggest a differential HLA-DRB1 expression depending on the allele. For example, one study showed consistently increased expression of *04:01 compared to *03:01 and *15:01 in different cell types of individuals with RA and healthy controls (30), whereas another publication on PBMCs of healthy donors and individuals with RA respectively suggest that this might be the case also for other alleles (31, 32). Taken together, these findings suggest differential expressions of alleles depending on the HLA-DRB1 genotype. However, larger studies are required for clarification, considering different cell types, tissues, allele-combinations, and other potentially cofounding factors (e.g., smoking). Optimally, these studies should investigate both the immunopeptidomics and transcriptomics level.

The fact that we found more peptides predicted to bind to SE alleles and *04:07 might explain, why more peptides were found in the RA groups compared to controls (Figure 1) as the RA groups contains more SE alleles (Table 1). In addition, it could explain why the presence of *04:01 and *04:05 genotypes were more indicative for sample similarity than for example *03:01 genotype (Figure 2) as the former were predicted to bind more identified peptides than the latter.

### 3.4 Comparison of the HLA-DR immunopeptidome between RA vs control and SE vs non-SE

Next, the dataset was searched for peptides which are specific for the RA group versus the control group using the Chi-square test (data not shown). Even though individual peptides (e.g. belonging to calreticulin) were found in 7/9 subjects with RA and none of the healthy controls, these results didn’t hold when investigating other peptides from the same nested set, i.e. sharing the same binding core. In these cases, it became apparent that the occurrence of these peptides in the RA group were related to the higher number of HLA-DRB1*04:01-positive individuals in the RA group and some peptides sharing the same binding core were also detected in the healthy control subjects which carried this genotype. In addition to this sampling bias, the fact that more peptides were detected in the RA group (Figure 1 H) further challenges this type of analysis. However, this does not prove the absence of RA-specific HLA-DR antigens in the lung either: Potential trigger-antigens might be present below the limit of detection or might be presented only transiently before the onset of clinical RA symptoms.

The dataset was also investigated for peptides which universally and selectively bind to SE alleles. Even though two peptides (from PRDX4 and LRP1) were assigned to bind to *01:01, *04:01, *04:05, and *10:01 by MHCMotifDecon in at least one sample per allele, and is also predicted to bind to *04:04 by NetMHCIIpan, this type of analysis is potentially biased: As more peptides have been predicted to bind to SE (see Figure 3), a peptide binding to SE is detected with a higher likelihood compared to a non-SE allele. Larger genotype-matched studies with higher availability of BAL cells are necessary to investigate this further with sufficient statistical power. However, considering how similar the binding preferences of some SE alleles are to non-SE alleles (e.g., *04:08 and *04:07) and how diverse the binding preference of most SE are, a SE-restricted universal antigen is not likely to be responsible for triggering RA. This must be considered especially since sero-positive RA can also develop in individuals without HLA-DR SE.

### 3.5 Known and potential RA antigens in the HLA-DR immunopeptidome of BAL cells

Autoantibodies against citrullinated proteins are specific for RA, however their role in the pathogenesis remains unknown. Immunogenic citrullinated self-peptides in RA patients presented on HLA-DR SE have been previously identified, suggesting that autoreactivity against citrullinated proteins does not only play a role on a B cell level but also in the immunological synapse between HLA-DR SE and the TCR (33–36).

However, in this dataset no citrullinated peptides could be identified. This is probably due to the scarce material, as 10^9^ cells per sample have been recommended for the study of post translational modifications in immunopeptidomics (37). Furthermore, the identification of citrullination poses methodical challenges in mass spectrometry, as the masses of citrulline and arginine differ by approximately 1 Dalton (38). Therefore, high quality mass spectra are required for a confident distinction, which in turn requires more cellular material. In order to narrow down the dataset for potential citrullinated autoepitopes for subsequent validation experiments, peptides containing arginine were extracted, which are potential citrullination sites (Table 2).

**Table 2:** Frequency of Arginine in relation to the HLA-DR binding pocket. . Number of peptides and binding cores predicted to bind to HLA-DR SE alleles, and number of binding cores with arginine in positions -2 to 9 in the HLA-DR-peptide binding cleft. Binding positions are highlighted in green

| SE allele | binding peptides | binding cores | Binding cores with arginine in |  |  |  |  |  |  |  |  |  |  |
| --- | --- | --- | --- | --- | --- | --- | --- | --- | --- | --- | --- | --- | --- |
|  |  |  | P-2 | P-1 | P1 | P2 | P3 | P4 | P5 | P6 | P7 | P8 | P9 |
| DRB1*01:01 | 2962 | 1039 | 70 | 56 | 0 | 91 | 49 | 0 | 82 | 0 | 60 | 94 | 2 |
| DRB1*04:01 | 3896 | 1509 | 101 | 85 | 1 | 135 | 77 | 0 | 82 | 0 | 23 | 129 | 11 |
| DRB1*04:04 | 1161 | 503 | 38 | 29 | 0 | 37 | 19 | 0 | 19 | 0 | 1 | 43 | 0 |
| DRB1*04:05 | 1905 | 802 | 45 | 45 | 0 | 55 | 31 | 0 | 24 | 0 | 11 | 68 | 3 |
| DRB1*10:01 | 2332 | 919 | 65 | 49 | 1 | 82 | 52 | 0 | 70 | 0 | 12 | 86 | 5 |

Previous immunogenicity studies on citrullinated peptides in RA showed that citrulline can invoke immunogenicity compared to the native peptides in two distinct manners: Citrulline in binding positions of the HLA-DR-peptide complex (e.g. in pocket 4 of *04:01 and *04:04) can modify the binding affinity of the peptide (39). Conversely, citrulline in non-binding positions can modulate the peptide-TCR interaction, e.g. P-2, P2 (33, 36). In Table 2, almost no peptides show arginine in the binding positions P1, P4, P6, and P9. This is expected as none of these SE alleles prefer basic amino acids in the binding positions, which is why the native counterparts of citrullinated peptides with citrulline in these positions would not necessarily show up in this analysis.

In non-binding positions, this analysis revealed peptides which would be of interest for further immunogenicity studies. As a proof of concept, we identified native peptides with already known immunogenicity for their citrullinated versions in RA: For example, we could identify 10 peptides from α-enolase sharing the sequence FRAAVPSGA which was predicted to bind to *01:01, *04:01, and *10:01. When citrullinated, this epitope has shown T-cell activation in a *04:01 restricted manner in a previous study (33). Also, 20 peptides from vimentin sharing the sequence VYATRSSA were identified, some of which are predicted to bind to *01:01 and *04:01. The citrullinated version has also been previously shown to elicit T cell response in RA (35). Fibrinogen β epitope GYRARPAKAAAT was detected, which has been previously shown to elicit T cell response against the citrullinated form in *04:01 (35). Also, 45 peptides belonging to known citrullinated RA target calreticulin were identified; however, we could not identify the T cell epitope which has been reported for RA in bronchiectasis (34).

Searches of the proteome of common lung microbiome as well as *porphyromonas gingivalis,* which has been previously implicated in RA, did not yield any peptide identifications (40). As discussed earlier, this does not signify absence of these epitopes as they might be presented transiently during pathogenesis or might be present below the limit of detection.

Finally, peptides containing known citrullination sites in the lung have been identified, belonging to ACTG1, EEF1A1, KRT17, PKM, and PPIA (38). These results show that tissue specific immunopeptidomics are a powerful tool to narrow down the proteome to potential T cell epitopes which can then be investigated in activation assays.

## 4. Conclusions

This study maps the HLA-DR immunopeptidome of the cells from bronchoalveolar lavage in subjects with RA and healthy controls with multiple HLA-DR genotypes. No significant differences between the RA and healthy groups could be observed, which were not confounded by the HLA-DRB1 genotype. However, motif deconvolution revealed that more peptides are predicted to bind to some HLA-DRB1 alleles, most of which belong to the SE risk alleles for RA. This is likely caused by allele-specific expression patterns but requires further studies. In the future, the peptides identified in this study will be screened for immunogenicity in T cell activation assays, which might give insight into the pathogenesis of RA.

## 5. Acknowledgements

We are grateful to the late Anca Catrina for her visionary role in conceptualizing this study. We also thank Monika Hansson for performing the ACPA testing, Rikard Holmdal for providing the antibody used in immunopurification, research nurses Helene Blomquist, Margitha Dahl and Gunnel DeForst for assistance during BAL and sample collection, biomedical technician Benita Engvall for assistance with sample work-up, and Peter Van Veelen for providing training in the immunopeptidomics workflow via the ArthritisHeal Marie Slodowska Curie doctoral training grant. These studies were funded by the EU (ArthritisHeal), the Swedish Heart-Lung Foundation, King Gustav V foundation, and the Swedish Research Council.

## References

1. Aletaha D, Neogi T, Silman AJ, Funovits J, Felson DT, Bingham CO, 3rd, et al. 2010 rheumatoid arthritis classification criteria: an American College of Rheumatology/European League Against Rheumatism collaborative initiative. Ann Rheum Dis. 2010;69(9):1580–8.

2. Collaborators GBDRA. Global, regional, and national burden of rheumatoid arthritis, 1990-2020, and projections to 2050: a systematic analysis of the Global Burden of Disease Study 2021. Lancet Rheumatol. 2023;5(10):e594–e610.

3. Ronnelid J, Wick MC, Lampa J, Lindblad S, Nordmark B, Klareskog L, et al. Longitudinal analysis of citrullinated protein/peptide antibodies (anti-CP) during 5 year follow up in early rheumatoid arthritis: anti-CP status predicts worse disease activity and greater radiological progression. Ann Rheum Dis. 2005;64(12):1744–9.

4. Padyukov L, Seielstad M, Ong RT, Ding B, Ronnelid J, Seddighzadeh M, et al. A genome-wide association study suggests contrasting associations in ACPA-positive versus ACPA-negative rheumatoid arthritis. Ann Rheum Dis. 2011;70(2):259–65.

5. Rantapaa-Dahlqvist S, de Jong BA, Berglin E, Hallmans G, Wadell G, Stenlund H, et al. Antibodies against cyclic citrullinated peptide and IgA rheumatoid factor predict the development of rheumatoid arthritis. Arthritis Rheum. 2003;48(10):2741–9.

6. Nielen MM, van Schaardenburg D, Reesink HW, van de Stadt RJ, van der Horst-Bruinsma IE, de Koning MH, et al. Specific autoantibodies precede the symptoms of rheumatoid arthritis: a study of serial measurements in blood donors. Arthritis Rheum. 2004;50(2):380–6.

7. Klareskog L, Stolt P, Lundberg K, Kallberg H, Bengtsson C, Grunewald J, et al. A new model for an etiology of rheumatoid arthritis: smoking may trigger HLA-DR (shared epitope)-restricted immune reactions to autoantigens modified by citrullination. Arthritis Rheum. 2006;54(1):38–46.

8. Reynisdottir G, Karimi R, Joshua V, Olsen H, Hensvold AH, Harju A, et al. Structural changes and antibody enrichment in the lungs are early features of anti-citrullinated protein antibody-positive rheumatoid arthritis. Arthritis Rheumatol. 2014;66(1):31–9.

9. Joshua V, Loberg Haarhaus M, Hensvold A, Wahamaa H, Gerstner C, Hansson M, et al. Rheumatoid Arthritis-Specific Autoimmunity in the Lung Before and at the Onset of Disease. Arthritis Rheumatol. 2023.

10. Pianta A, Chiumento G, Ramsden K, Wang Q, Strle K, Arvikar S, et al. Identification of Novel, Immunogenic HLA-DR-Presented Prevotella copri Peptides in Patients With Rheumatoid Arthritis. Arthritis Rheumatol. 2021;73(12):2200–5.

11. Wang Q, Drouin EE, Yao C, Zhang J, Huang Y, Leon DR, et al. Immunogenic HLA-DR-Presented Self-Peptides Identified Directly from Clinical Samples of Synovial Tissue, Synovial Fluid, or Peripheral Blood in Patients with Rheumatoid Arthritis or Lyme Arthritis. J Proteome Res. 2017;16(1):122–36.

12. Maggi J, Carrascal M, Soto L, Neira O, Cuéllar MC, Aravena O, et al. Isolation of HLA-DR-naturally presented peptides identifies T-cell epitopes for rheumatoid arthritis. Annals of the Rheumatic Diseases. 2022;81(8):1096–105.

13. Seward RJ, Drouin EE, Steere AC, Costello CE. Peptides presented by HLA-DR molecules in synovia of patients with rheumatoid arthritis or antibiotic-refractory Lyme arthritis. Mol Cell Proteomics. 2011;10(3):M110 002477.

14. Kaabinejadian S, Barra C, Alvarez B, Yari H, Hildebrand WH, Nielsen M. Accurate MHC Motif Deconvolution of Immunopeptidomics Data Reveals a Significant Contribution of DRB3, 4 and 5 to the Total DR Immunopeptidome. Front Immunol. 2022;13:835454.

15. Kohler M, Sandberg A, Kjellqvist S, Thomas A, Karimi R, Nyren S, et al. Gender differences in the bronchoalveolar lavage cell proteome of patients with chronic obstructive pulmonary disease. J Allergy Clin Immunol. 2013;131(3):743–51.

16. Eklund A, Blaschke E. Relationship between changed alveolar-capillary permeability and angiotensin converting enzyme activity in serum in sarcoidosis. Thorax. 1986;41(8):629–34.

17. Lofdahl JM, Cederlund K, Nathell L, Eklund A, Skold CM. Bronchoalveolar lavage in COPD: fluid recovery correlates with the degree of emphysema. Eur Respir J. 2005;25(2):275–81.

18. Teo GC, Polasky DA, Yu F, Nesvizhskii AI. Fast Deisotoping Algorithm and Its Implementation in the MSFragger Search Engine. J Proteome Res. 2021;20(1):498–505.

19. Yang KL, Yu F, Teo GC, Li K, Demichev V, Ralser M, et al. MSBooster: improving peptide identification rates using deep learning-based features. Nat Commun. 2023;14(1):4539.

20. Kong AT, Leprevost FV, Avtonomov DM, Mellacheruvu D, Nesvizhskii AI. MSFragger: ultrafast and comprehensive peptide identification in mass spectrometry-based proteomics. Nat Methods. 2017;14(5):513–20.

21. Kall L, Canterbury JD, Weston J, Noble WS, MacCoss MJ. Semi-supervised learning for peptide identification from shotgun proteomics datasets. Nat Methods. 2007;4(11):923–5.

22. Nesvizhskii AI, Keller A, Kolker E, Aebersold R. A statistical model for identifying proteins by tandem mass spectrometry. Anal Chem. 2003;75(17):4646–58.

23. da Veiga Leprevost F, Haynes SE, Avtonomov DM, Chang HY, Shanmugam AK, Mellacheruvu D, et al. Philosopher: a versatile toolkit for shotgun proteomics data analysis. Nat Methods. 2020;17(9):869–70.

24. Zhang L, McAlpine PL, Heberling ML, Elias JE. Automated Ligand Purification Platform Accelerates Immunopeptidome Analysis by Mass Spectrometry. J Proteome Res. 2021;20(1):393–408.

25. Ikeda N, Kojima H, Nishikawa M, Hayashi K, Futagami T, Tsujino T, et al. Determination of HLA-A, -C, -B, -DRB1 allele and haplotype frequency in Japanese population based on family study. Tissue Antigens. 2015;85(4):252–9.

26. Creary LE, Sacchi N, Mazzocco M, Morris GP, Montero-Martin G, Chong W, et al. High-resolution HLA allele and haplotype frequencies in several unrelated populations determined by next generation sequencing: 17th International HLA and Immunogenetics Workshop joint report. Hum Immunol. 2021;82(7):505–22.

27. Wang J, Jelcic I, Muhlenbruch L, Haunerdinger V, Toussaint NC, Zhao Y, et al. HLA-DR15 Molecules Jointly Shape an Autoreactive T Cell Repertoire in Multiple Sclerosis. Cell. 2020;183(5):1264–81 e20.

28. Heyder T, Kohler M, Tarasova NK, Haag S, Rutishauser D, Rivera NV, et al. Approach for Identifying Human Leukocyte Antigen (HLA)-DR Bound Peptides from Scarce Clinical Samples. Mol Cell Proteomics. 2016;15(9):3017–29.

29. Chong C, Marino F, Pak H, Racle J, Daniel RT, Muller M, et al. High-throughput and Sensitive Immunopeptidomics Platform Reveals Profound Interferongamma-Mediated Remodeling of the Human Leukocyte Antigen (HLA) Ligandome. Mol Cell Proteomics. 2018;17(3):533–48.

30. Houtman M, Hesselberg E, Ronnblom L, Klareskog L, Malmstrom V, Padyukov L. Haplotype-Specific Expression Analysis of MHC Class II Genes in Healthy Individuals and Rheumatoid Arthritis Patients. Front Immunol. 2021;12:707217.

31. Chun S, Bang SY, Ha E, Cui J, Gu KN, Lee HS, et al. Allele-Specific Quantification of HLA–DRB1 Transcripts Reveals Imbalanced Allelic Expression That Modifies the Amino Acid Effects in HLA–DRβ1. Arthritis & Rheumatology. 2021;73(3):381–91.

32. Yamamoto F, Suzuki S, Mizutani A, Shigenari A, Ito S, Kametani Y, et al. Capturing Differential Allele-Level Expression and Genotypes of All Classical HLA Loci and Haplotypes by a New Capture RNA-Seq Method. Front Immunol. 2020;11:941.

33. Gerstner C, Dubnovitsky A, Sandin C, Kozhukh G, Uchtenhagen H, James EA, et al. Functional and Structural Characterization of a Novel HLA-DRB1*04:01-Restricted alpha-Enolase T Cell Epitope in Rheumatoid Arthritis. Front Immunol. 2016;7:494.

34. Clarke A, Perry E, Kelly C, De Soyza A, Heesom K, Gold LI, et al. Heightened autoantibody immune response to citrullinated calreticulin in bronchiectasis: Implications for rheumatoid arthritis. Int J Biochem Cell Biol. 2017;89:199–206.

35. Gerstner C, Turcinov S, Hensvold AH, Chemin K, Uchtenhagen H, Ramwadhdoebe TH, et al. Multi-HLA class II tetramer analyses of citrulline-reactive T cells and early treatment response in rheumatoid arthritis. BMC Immunol. 2020;21(1):27.

36. Chemin K, Pollastro S, James E, Ge C, Albrecht I, Herrath J, et al. A Novel HLA-DRB1*10:01-Restricted T Cell Epitope From Citrullinated Type II Collagen Relevant to Rheumatoid Arthritis. Arthritis Rheumatol. 2016;68(5):1124–35.

37. Stutzmann C, Peng J, Wu Z, Savoie C, Sirois I, Thibault P, et al. Unlocking the potential of microfluidics in mass spectrometry-based immunopeptidomics for tumor antigen discovery. Cell Rep Methods. 2023;3(6):100511.

38. Lee CY, Wang D, Wilhelm M, Zolg DP, Schmidt T, Schnatbaum K, et al. Mining the Human Tissue Proteome for Protein Citrullination. Mol Cell Proteomics. 2018;17(7):1378–91.

39. Scally SW, Petersen J, Law SC, Dudek NL, Nel HJ, Loh KL, et al. A molecular basis for the association of the HLA-DRB1 locus, citrullination, and rheumatoid arthritis. J Exp Med. 2013;210(12):2569–82.

40. Sherina N, de Vries C, Kharlamova N, Sippl N, Jiang X, Brynedal B, et al. Antibodies to a Citrullinated Porphyromonas gingivalis Epitope Are Increased in Early Rheumatoid Arthritis, and Can Be Produced by Gingival Tissue B Cells: Implications for a Bacterial Origin in RA Etiology. Front Immunol. 2022;13:804822.

